# Metabarcoding for the parallel identification of several hundred predators and their preys: application to bat species diet analysis

**DOI:** 10.1101/155721

**Authors:** Maxime Galan, Jean-Baptiste Pons, Orianne Tournayre, Éric Pierre, Maxime Leuchtmann, Dominique Pontier, Nathalie Charbonnel

## Abstract

Assessing diet variability is of main importance to better understand the biology of bats and design conservation strategies. Although the advent of metabarcoding has facilitated such analyses, this approach does not come without challenges. Biases may occur throughout the whole experiment, from fieldwork to biostatistics, resulting in the detection of false negatives, false positives or low taxonomic resolution. We detail a rigorous metabarcoding approach based on a short COI minibarcode and two-step PCR protocol enabling the ‘all at once’ taxonomic identification of bats and their arthropod preys for several hundreds of samples. Our study includes faecal pellets collected in France from 357 bats representing 16 species, as well as insect mock communities that mimic bat meals of known composition, negative and positive controls. All samples were analysed using three replicates. We compare the efficiency of DNA extraction methods and we evaluate the effectiveness of our protocol using identification success, taxonomic resolution, sensitivity, and amplification biases. Our parallel identification strategy of predators and preys reduces the risk of mis-assigning preys to wrong predators and decreases the number of molecular steps. Controls and replicates enable to filter the data and limit the risk of false positives, hence guaranteeing high confidence results for both prey occurrence and bat species identification. We validate 551 COI variants from arthropod including 18 orders, 117 family, 282 genus and 290 species. Our method therefore provides a rapid, resolutive and cost-effective screening tool for addressing evolutionary ecological issues or developing ‘chirosurveillance’ and conservation strategies.

## Introduction

DNA metabarcoding has revolutionized our approaches of biodiversity assessment this last decade (Taberlet, Coissac, Pompanon, Brochmann, & Willerslev, 2012). The method is based on the high-throughput sequencing (HTS) of DNA barcode regions (i.e. small fragments of DNA that exhibit low intra-species and high inter-species variability), which are amplified using universal PCR primers (Hebert, Cywinska, Ball, & deWaard, 2003). Nowadays recent HTS technologies generate millions of sequences concurrently. Metabarcoding therefore enables to characterize quickly and in a single experiment a very large number of species present in an environmental sample, and also to analyse simultaneously several hundreds of samples by using tags / index and multiplexing protocols (Binladen et al., 2007). Metabarcoding has proved to be useful for a wide variety of applications (Bohmann et al., 2014; Taberlet et al., 2012).

Dietary analyses have been facilitated by the advent of metabarcoding and its application to the analysis of faeces or stomach contents (Pompanon et al., 2012). Compared to traditional morphological analyses of remaining hard parts, a large sampling can be processed quickly. Results are more sensitive, allowing the identification of a larger array of specimens (e.g. juvenile life stages) and providing a greater taxonomic resolution (e.g. cryptic species might be detected, Hebert, Penton, Burns, Janzen, & Hallwachs, 2004). In addition, traditional molecular approaches (real time PCRs, Sanger sequencing) require several assays or reactions to discriminate the different taxa present in the same sample, while a single run of metabarcoding provides the identification of a broad range of taxa with no *a priori* (Tillmar, Dell’Amico, Welander, & Holmlund 2013). Research on insectivorous bat dietary analyses have been pioneering in benefiting from these advantages of DNA metabarcoding (Bohmann et al., 2011). Indeed, direct observations of prey consumption are made difficult as these species fly and are nocturnal. Moreover, because many bat species are vulnerable or endangered around the world, catching might be difficult, even forbidden during hibernation, and invasive methods cannot be applied. Morphologic examinations of faeces and guano have therefore initially provided important knowledge on bat diets (Hope et al., 2014; Lam et al., 2013). These methods have major limits (time consuming, taxonomic expertise required, low resolution and ascertainment biases due to the reject of insect hard parts, see refs in Iwanowicz et al., 2016). In particular, identifying preys at the species level is not possible based on morphological analyses of faecal samples.

Obtaining reliable and reproducible results from metabarcoding is not straightforward. Several biases may occur throughout the whole experiment, resulting in the detection of false negatives, false positives or low taxonomic resolution (Ficetola, Taberlet, & Coissac, 2016). These biases take place from fieldwork to biostatistics (see for a review in an epidemiological context, Galan et al., 2016). Contaminations occurring during sampling or in the laboratory (Champlot et al., 2010; Goldberg et al., 2016) may be further amplified and sequenced due to the high sensitivity of the PCR and to the high depth of sequencing provided by the HTS, leading to further misinterpretations (see examples in Ficetola et al., 2016). The choice of the DNA extraction method, the barcode region and primers may also influence the issue of metabarcoding studies. Low efficiencies of sample disruption, high losses of genetic material or the presence of PCR inhibitors may lead to false negative results (Deiner, Walser, Machler, & Altermatt, 2015). Incorrect design of primers and unsuitable barcode region may prevent or bias the amplification of the taxonomic taxa studied, or may result in an identification at low resolution levels (Hajibabaei et al., 2006). In addition to these well-known precautions required for metabarcoding studies, other considerations need to be made. First, multiplexing samples within HTS runs results in mis-assignments of reads after bioinformatic demultiplexing (Kircher, Sawyer, & Meyer, 2012). The detection of one or a few reads assigned to a given taxon in a sample therefore does not necessarily mean this taxon is actually present in that sample (Galan et al., 2016). These errors may originate from: (i) contamination of tags/index, (ii) production of between samples chimera due to jumping PCR (Schnell, Bohmann, & Gilbert, 2015) when libraries require the bulk amplification of tagged samples, or (iii) the presence of mixed cluster of sequences (i.e. polyclonal clusters) on the Illumina flowcell surface. They may have dramatic consequences on the proportion of false positives (e.g. up to 28.2% of mis-assigned unique sequences reported in Esling, Lejzerowicz, & Pawlowski, 2015). Unfortunately, read mis-assignments due to polyclonal clusters during Illumina sequencing are difficult to avoid and concern 0.2 to 0.6% of the reads generated (Galan et al., 2016; Kircher et al., 2012; Wright & Vetsigian, 2016). It is therefore of main importance to filter the occurrence data obtained through metabarcoding experiments, using both controls and replicates (Ficetola et al., 2016; Galan et al., 2016; Robasky, Lewis, & Church, 2014). The second set of parameters still scarcely considered during metabarcoding experiments includes the sensitivity and taxonomic resolution of the protocol designed. They can be assessed empirically by analysing mock communities (MC), i.e. pools of DNA belonging to different species, hence simulating a predator meal of known composition (Pinol, Mir, Gomez-Polo, & Agusti, 2015).

Here, we propose a rigorous metabarcoding approach based on a two-step PCR protocol and bioinformatic analyses enabling the ‘*all at once’* identification, potentially at the species level, of bats and their arthropod preys for several hundreds of samples. We use faecal pellets from 357 bats representing 16 species. Our aims are threefold. First, we compare the efficiency of DNA extraction and purification methods among six available commercial kits. Second, we design a scrupulous experimental protocol that includes negative and positive controls as well as systematic technical replicates. They enable to filter occurrence results (Ficetola et al., 2016; Galan et al., 2016). Then, we evaluate the effectiveness of this protocol using a set of criteria including the rate of identification success, taxonomic resolution, sensitivity, and amplification biases. To this end, we analyse arthropod mock communities and we validate bat identifications by comparing molecular results with morphological identifications performed during fieldwork by experts. Third, we apply this DNA metabarcoding protocol to identify the consumed preys of the 357 faecal bat samples analysed, and we examine our results with regard to dietary analyses of bats previously published in the literature.

## Materials and methods

### Study sites and sample collection

Bats were captured from summer roost sites between June and September 2015 in 18 sites located in Western France (Poitou-Charentes). For each site, harp traps were placed one night, at the opening of cave or building before sunset. Each captured bat was placed in a cotton holding bag until it was weighted, sexed and measured. Species identification was determined based on morphological criteria. Bats were then released. All faecal pellets were collected from holding bag, and stored in microtubes at room temperature until DNA was extracted 45 to 162 days later. Storage conditions did not follow the recommendations described for metabarcoding studies, as samples were initially collected for diet analyses based on morphological identifications. Authorization for bat capture was provided by the Ministry of Ecology, Environment, and Sustainable development over the period 2015-2020 (approval no. C692660703 from the Departmental Direction of Population Protection (DDPP, Rhône, France). All methods were approved by the MNHN and the SFEPM.

### Laboratory precautions and controls

Throughout the experiment, we strictly applied the laboratory protocols to prevent contamination by alien DNA and PCR products. All pre-PCR laboratory manipulations were conducted with filter tips under a sterile hood in a DNA-free room. The putative presence of contamination was checked at this stage and along the whole laboratory procedure using different negative and positive controls. A large number of research has highlighted several biases occurring at different steps of amplicon HTS (for a detailed list, see Galan et al., 2016). These biases can be estimated directly from data by including several controls together with samples in the experiment (for details see Appendix S1): negative controls for DNA extraction (NC_ext_), negative controls for PCR (NC_PCR_), negative controls for indexing (NC_index_: unused dual-index combinations), positive controls for PCR (PC_PCR_) including mock communities (MC) and positive controls for indexing (PC_alien_: DNA from beluga whale -*Delphinapterus leucas*- used to estimate the read mis-assignment frequency).

### DNA extraction from faecal samples

We analysed faecal pellets from 357 bats corresponding to 16 species. Details are provided in Table S1 (Supplemental Information). One pellet per individual was frozen at −80°C, bead-beaten for 2 x 30s at 30Hz on a TissueLyser (Qiagen) using a 5mm stainless steel bead then extracted. We randomised the 357 faecal samples between six silica-membrane DNA extraction kits to compare their efficiency: EZ-10 96 DNA Kit, Animal Samples (BioBasic; *n* = 113), QIAamp Fast DNA Stool Mini Kit (Qiagen; *n* = 47), DNeasy mericon Food Kit (Qiagen; *n* = 47), ZR Fecal DNA MiniPrep (Zymo; *n* = 46), NucleoSpin 8 Plant II (Macherey-Nagel; *n* = 95) and NucleoSpin Soil (Macherey-Nagel; *n* = 9). The EZ-10 96 DNA and NucleoSpin 8 Plant II kits provide high-throughput DNA isolation (up to 192 samples in parallel) thanks to a 96-well format unlike the other kits using tube format. For all kits, we followed manufacturer's recommendations except for the NucleoSpin 8 Plant II as we used the slight modifications recommended in Zarzoso-Lacoste et al. (2017).

We compared DNA extraction kits' efficiency using three criteria: i) the mean number of reads per PCR obtained after sequencing; ii) the success rate of host sequencing (presence of chiropter reads from the same variant, found repeatedly between the three PCR replicates) and iii) the success rate of prey sequencing (presence of variants corresponding to arthropods, found repeatedly between the three PCR replicates). Because storage duration could have influenced sequencing results, we included this variable in the statistical models performed. The number of reads was analysed with a Gaussian function and the success of sequencing was analysed with a binomial function and a logit error (see Appendix S2 for details of statistical analyses). All analyses were performed in R 3.1.0 (R core team, 2013).

### Mock community preparation

To better evaluate the sensitivity of our metabarcoding approach, we created two artificial communities of arthropods that mimic insectivorous bat diets. The first mock community (MC_1_) was composed of 12 species and the second one (MC_2_) included seven species (see details in Table 1 and Table S1).

**Table 1.**
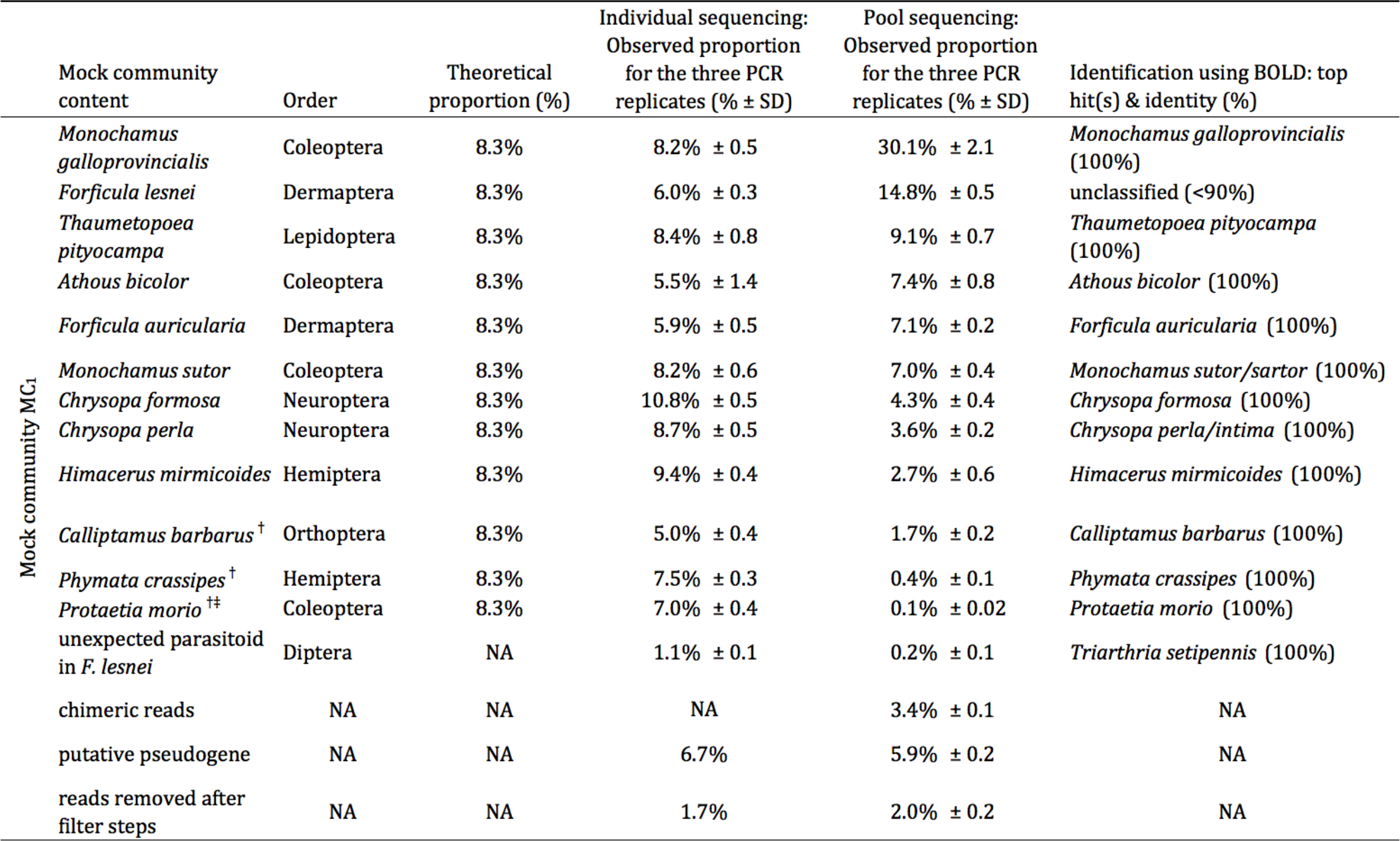
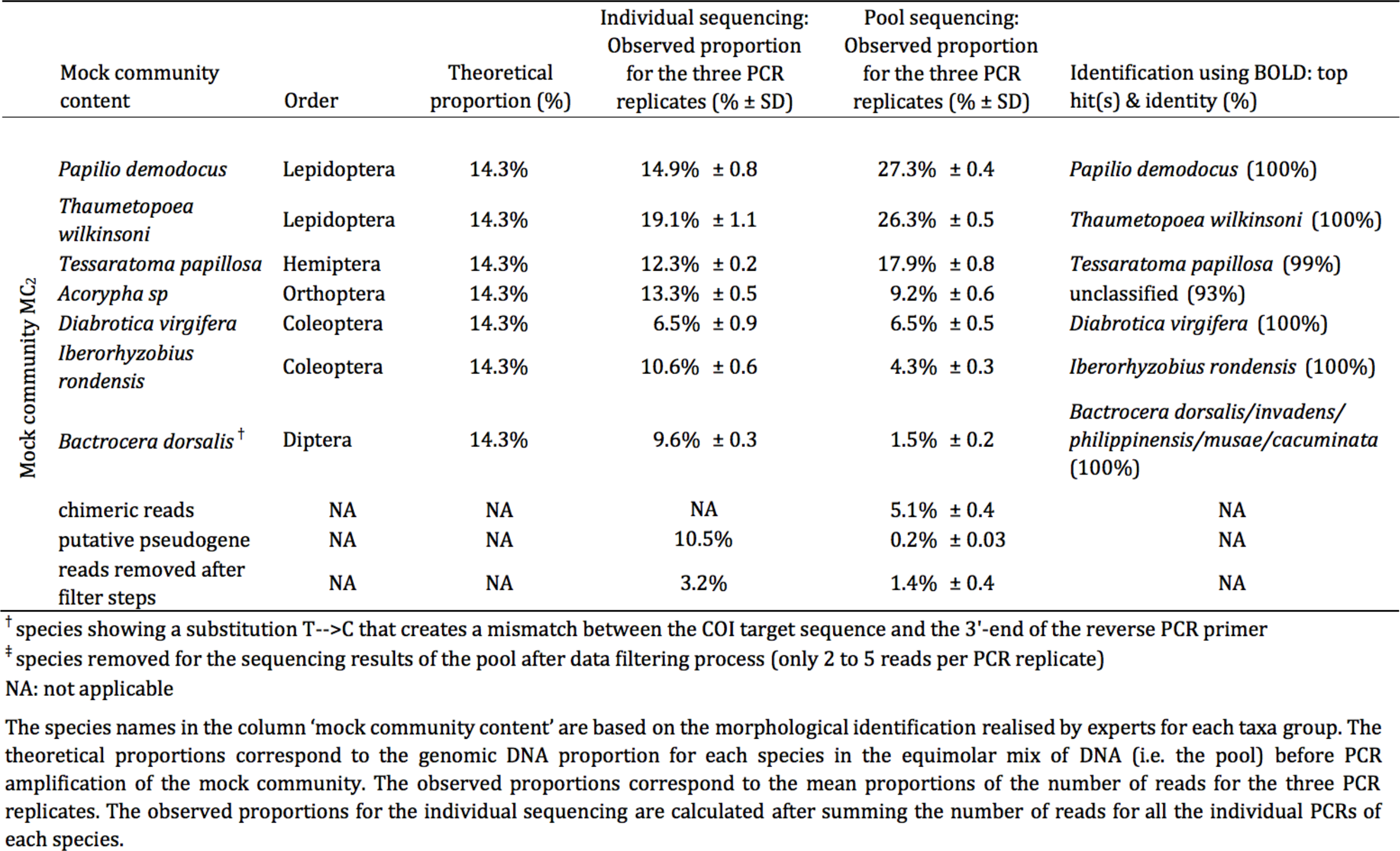
Proportion of reads for two arthropod mock communites (MC_1_ & MC_2_)

Arthropod DNA was extracted individually using the DNeasy Blood & Tissue kit (Qiagen). Sanger sequences were available for the cytochrome oxydase I (COI) gene of each individual (see the alignment file of the 19 insect species included in the mock communities deposited in Dryad: https://datadryad.org/resource/doi:10.5061/dryad.kv02g/8Dryad). DNA extractions were normalized to 5ng/µL using Qubit fluorimeter quantification (Invitrogen). First, each normalized DNA was amplified and sequenced independently. Second, normalized DNA extractions were pooled in equal proportion to build the two mock communities MC_1_ and MC_2._ These latter were amplified and sequenced (see details in Table S1). Results provided by independent (insect individual) and pooled (mock communities) sequencing were compared. It enabled to estimate biases resulting from the co-amplification of different species mixed within the same reaction.

### COI minibarcode, PCR and library construction

We used the 133 bp minibarcode of COI described in Gillet et al. (2015). Its efficiency to identify a wide taxonomic range of arthropods from France has been proven recently: 20 arthropod orders were detected in the diet of *Galemys pyrenaicus* and 24 in the diet of *Neomys fodiens* (Biffi, Gillet, et al., 2017; Biffi, Laffaille, et al., 2017). We have verified the discriminatory power of this minibarcode for resolving bat species identification using an *in silico* analysis based on 444 BOLD (Barcode of Life DataBase) reference sequences corresponding to the 33 bat species found in France (see the phylogenetic tree provided in Figure S1 and the Table S2).

We performed a two-step PCR strategy (see Illumina Application Note Part 15044223) combined with the dual-index paired-end sequencing approach described in Kozich et al. 2013 (Fig. 1). The 32 index i5 and 48 index i7 allow to multiplex up to 1536 PCR products in the same MiSeq run. This makes it possible to multiplex several hundreds of samples while performing several technical replicates per sample. The two-step PCR strategy enables to build all libraries independently for each sample and technical replicate, without pre-sequencing PCR enrichment of a mix of PCR products from different samples. This method prevents the production of between-samples chimera due to jumping PCR (Schnell et al., 2015). It therefore reduces drastically the risk of mistaging/misindexing.

**Figure 1.**
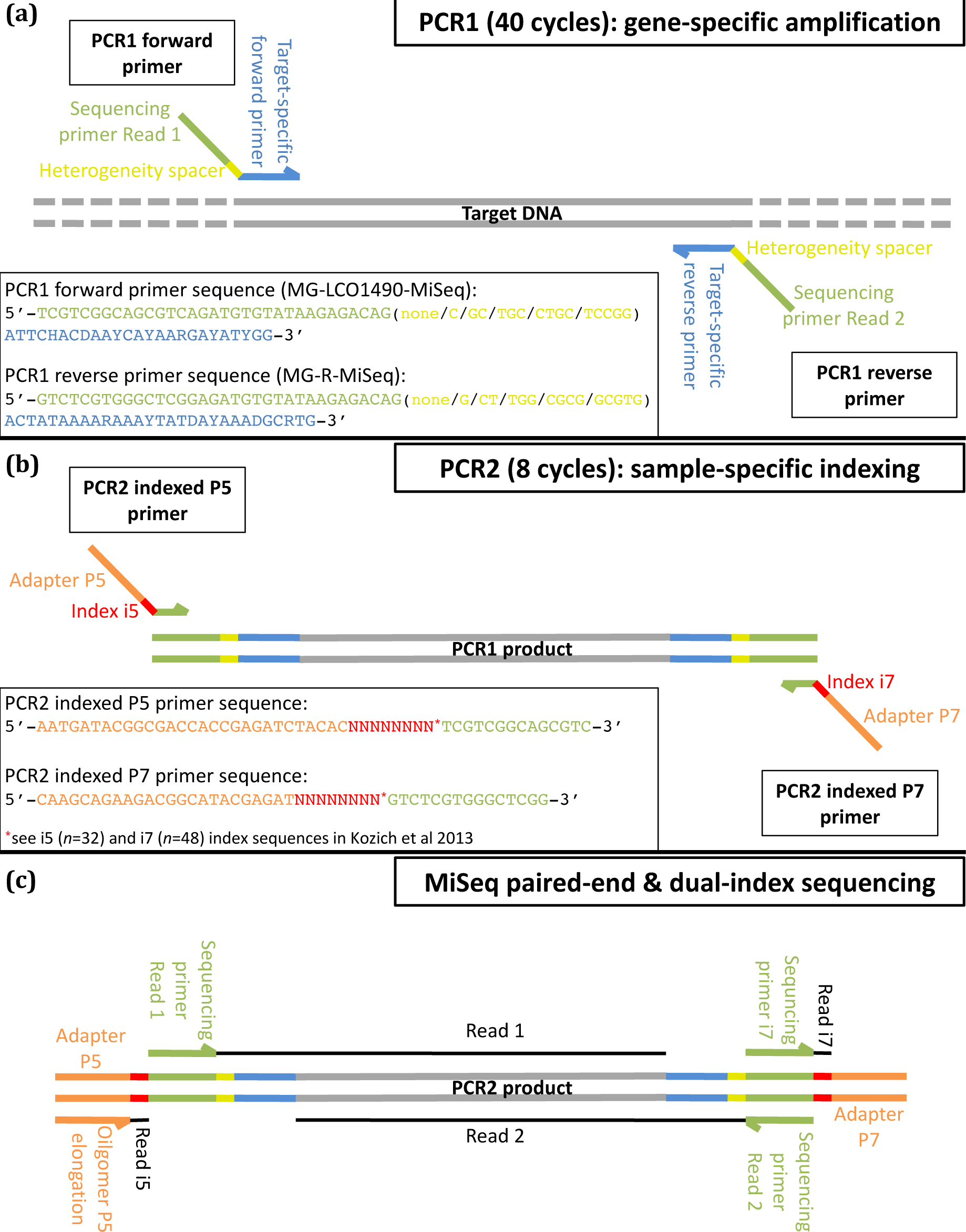
Schematic description of the library construction using 2-step PCR and MiSeq sequencing. (a) During PCR_1_, the COI minibarcode was amplified for each sample using the gene specific forward and reverse primers (in blue) tailed with the overhang Illumina sequencing primer sequences (in green), and alternative bases called heterogeneity spacers (in yellow) to create an artificial nucleotide diversity during the first cycles of the Illumina sequencing. (b) The PCR_2_ aims at adding the Illumina adapters (in orange) and the multiplexing dual-indices (in red) to each sample replicate by performing a limited-cycle amplification step. (c) After PCR_2_, all sample libraries are pooled together and then sequenced. During the MiSeq paired-end sequencing, each nucleotide of the target gene is read twice (read 1 & read 2) and the dual-index reads allow to assign the millions of sequences to the corresponding original samples. For the future studies, we propose a new version of the target-specific reverse primer MG-univR: 5’-ACTATAAARAARATYATDAYRAADGCRTG-3’ (see details in Discussion)

During the first PCR (PCR_1_), we used a primer pair corresponding to highly modified and degenerated versions of forward primer LepF1 (Hebert et al., 2004) and reverse primer EPT-long-univR (Hajibabaei, Shokralla, Zhou, Singer, & Baird, 2011). We added the partial overhang Illumina sequencing primers in 5’-end and a heterogeneity spacer of each target-specific primer (Figure 1): MG-LCO1490-MiSeq 5’-TCGTCGGCAGCGTCAGATGTGTATAAGAGACAG(none/C/GC/TGC/CTGC/TCCGG) ATTCHACDAAYCAYAARGAYATYGG-3’ and MG-R-MiSeq 5’-GTCTCGTGGGCTCGGAGATGTGTATAAGAGACAG(none/G/CT/TGG/CGCG/GCGTG) ACTATAAAARAAAYTATDAYAAADGCRTG-3’. The alternative bases between the partial adaptors and the target-specific primers correspond to a 0 to 5 bp “heterogeneity spacer” designed to mitigate the issues caused by low sequence diversity in Illumina amplicon sequencing (Fadrosh et al., 2014). These five versions of each forward and reverse primer were mixed together before PCR_1_. During the first cycles of the Illumina sequencing, they created an artificial diversity of the four nucleobases to improve the detection of the sequencing clusters at the flowcell surface, what consequently increased the quality of the reads. This PCR_1_ was performed in 11 µL reaction volume using 5 µL of 2× Qiagen Multiplex Kit Master Mix (Qiagen), 0.5 µM of each mix of forward and reverse primers, and 2 µL of DNA extract. The PCR conditions consisted in an initial denaturation step at 95°C for 15 min, followed by 40 cycles of denaturation at 94°C for 30 s, annealing at 45°C for 45 s, and extension at 72°C for 30 s, followed by a final extension step at 72°C for 10 min.

During the second PCR (PCR_2_), we used the PCR_1_ as DNA template. This PCR_2_ consists of a limited-cycle amplification step to add multiplexing indices i5 and i7 and Illumina sequencing adapters P5 and P7 at both ends of each DNA fragment. The indexed primers P5 (5’-AATGATACGGCGACCACCGAGATCTACAC(8-base i5 index)TCGTCGGCAGCGTC-3’) and P7 (5’-CAAGCAGAAGACGGCATACGAGAT(8-base i7 index)GTCTCGTGGGCTCGG-3’) were synthetized with the different 8-base index sequences described in Kozich et al. (Kozich, Westcott, Baxter, Highlander, & Schloss, 2013). PCR_2_ was carried out in a 11 µL reaction volume using 5 µL of Qiagen Multiplex Kit Master Mix (Qiagen) and 0.7 µM of each indexed primer. Then, in a post-PCR room, 2 µL of PCR_1_ product was added to each well. The PCR_2_ started by an initial denaturation step of 95°C for 15 min, followed by 8 cycles of denaturation at 94°C for 40 s, annealing at 55°C for 45 s and extension at 72°C for 60 s followed by a final extension step at 72°C for 10 min.

PCR_2_ products (3 μL) were verified by electrophoresis in a 1.5% agarose gel. One PCR blank (NC_PCR_) and one negative control for indexing (NC_index_) were systematically added to each of the 15 PCR microplates. Each DNA extraction was amplified and indexed in three independent PCR reactions. These PCR replicates were used as technical replicates to confirm the presence of a taxa in a sample and further remove the false positive results (Robasky et al., 2014). A MiSeq (Illumina) run was conducted, including PCR_2_ products from bat faecal samples (number of PCRs *n* = 357 x 3), the positive (*n* = 21 x 3 arthropods used in the mock communities and *n* = 9 PCR replicates from a DNA beluga whale used as positive internal control PC_alien_) and negative (*n* = 58) controls (see details in Table S1).

### MiSeq sequencing

The MiSeq platform was chosen because it generates lower error rates than other HTS platforms (D’Amore et al., 2016). For this study, the number of PCR products multiplexed was 1168 (Table S1). PCR products were pooled by volume for each 96-well PCR microplate. Mixes were checked by electrophoresis in 1.5% agarose gel before generating a 'super-pool’ including all PCR products. We subjected 60 µL of the super-pool to size selection for the full-length amplicon (expected size: 312 bp including primers, indexes and adaptors) by excision on a low-melting agarose gel (1.25%). It enabled to discard non-specific PCR products and primer dimers. The PCR Clean-up Gel Extraction kit (Macherey-Nagel) was used to purify the excised band. The super-pool of amplicon libraries was quantified using the KAPA library quantification kit (KAPA Biosystems) before loading 8 pM and 10% of PhiX control on a MiSeq flow cell (expected cluster density: 700-800 K/mm2) with a 500-cycle Reagent Kit v2 (Illumina). We performed a run of 2 x 200 bp paired-end sequencing, which yielded high-quality sequencing through the reading of each nucleotide of the COI minibarcode fragments twice after the assembly of reads 1 and reads 2 (see details below). Information about PCR products and fastq file names are provided in Table S1.

### Sequence analyses and data filtering

We used MOTHUR program v1.34 (Schloss et al., 2009) to create an abundance table for each variant and each PCR product (Table S3). Briefly, MOTHUR enabled to i) contig the paired-end read 1 and read 2; ii) remove the reads with low quality of assembling (> 200 bp); iii) dereplicate the redundant reads; iv) align the variants on a COI minibarcode reference alignment; v) remove PCR primers; vi) remove misaligned variants; vii) correct a part of the PCR and sequencing errors by clustering the variants that are within one mutation of each other for each PCR replicate independently; viii) remove the singleton and rare variants (*cut-off =* 8 reads) and ix) remove chimeric variants using UCHIME (Edgar, Haas, Clemente, Quince, & Knight, 2011). Note that step i) enabled to remove an important number of sequencing errors: the pairs of sequences were aligned and any positions with discongruence between the two reads were identified and corrected using the quality score of each read position (see https://www.mothur.org/wiki/MiSeq_SOP). If one sequence exhibited a base and the other one a gap, the quality score of the base had to be higher than 25 to be considered real. If both sequences had a base at that given position, then we required one of the bases to have a quality score of six or more points than the other. Step vii) is applied independently for each sample and technical replicate by keeping only the most abundant variant among the cluster of similar variants at one mutation. This procedure enables to remove an important part of PCR or sequencing errors, and to validate variants differing by a single mutation when they are found in different PCR replicates or biological samples. Hereafter, a variant will correspond to a cluster of similar reads obtained for a given technical replicate and potentially differing by a single mutation difference.

We used the multiple controls introduced during the process to estimate potential biases and define read number thresholds above which the PCR product may be considered as positive for a given sequence. Following Galan et al. (2016), two different thresholds were set for each variant, and a cross-validation using the three PCR replicates was applied to confirm the positivity of each sample for each variant.

First, T_CC_ threshold was used to filter cross-contaminations during the laboratory procedure. For each variant, we used the maximum number of reads observed in the different negative controls (NC) as threshold. PCR products with fewer than this number of reads for this particular variant were not considered to be positive because this number of reads is not distinguishable from noise. For the positive samples, the information of the observed number of reads for each variant is kept. Second, T_FA_ threshold was applied to filter the false-assignments of reads to a PCR product due to the generation of mixed clusters during the sequencing (Kircher et al., 2012). This phenomenon was estimated in our experiment using “alien” positive controls (PC_alien_) sequenced in parallel with the bat faecal samples. As the PC_alien_ (i.e. DNA from beluga whale) were handled separately from bat samples before sequencing, the presence of reads from beluga whale in a bat sample indicated a sequence assignment error due to the Illumina sequencing (i.e. generation of mixed clusters). We determined the maximal number of reads of beluga whale assigned to a bat PCR product. We then calculated the false-assignment rate (R_FA_) for this PCR product by dividing this number of reads by the total number of reads from beluga in the sequencing run. Moreover, the number of reads for a specific variant mis-assigned to a PCR product should increase with the total number of reads of this variant in the MiSeq run. We therefore defined T_FA_ threshold as the total number of reads in the run for a variant multiplied by R_FA_. PCR products with fewer than T_FA_ for a particular variant were considered to be negative. We then discarded positive results associated with numbers of reads below the thresholds T_CC_ and T_FA_. Lastly, we discarded not-replicated positive results for the three PCR replicates to remove inconsistent variants due to PCR or sequencing errors or unconfident variants that could be associated with remaining false positive results. Finally, for each sample and variant, the reads obtained for the three PCR replicates were summed.

### Taxonomic assignment

We used BOLD Identification System (Ratnasingham & Heber, 2007) and species level barcode records (2,695,529 sequences/175,014 species in January 2017) to provide a taxonomic identification of each variant passing our filtering processes. We provided a 5-class criteria describing the confidence level of the sequence assignments, modified from Razgour et al. (2011). They were applied to hits with similarity higher than 97% (see details in Appendix S3). For multi-taxonomic affiliation, we kept the common parent of all possible taxa. For sequences exhibiting similarity results lower than 97% in BOLD, we performed a BLAST in GENBANK to improve the taxonomic identification. Finally, results with similarity lower than 97% in BOLD and GENBANK were assigned to the phylum level using the closest taxa, or were considered as unclassified taxa when no match was found in the databases.

## Results

### Sequencing results & data filtering

The MiSeq sequencing of 1162 PCR products including bat samples, positive and negative controls analysed in three PCR replicates generated a total paired-end read output of 6,918,534 reads of the COI minibarcode. MOTHUR program removed seven negative controls because they produced less than 8 reads, 633,788 (9.2%) of paired-end reads because they were misassembled, 312,336 (4.5%) of reads because they were misaligned, 125,606 (1.8%) of reads because they corresponded to rare variants (< 9 reads) and 11,445 (0.2%) of reads because they were chimeric (Table S3). The remaining reads represented a total of 5751 variants and 5,835,359 reads. The abundance table produced was next filtered.

#### Filtering cross-contaminations using threshold T_CC_

We observed between 0 and 6,230 reads (total: 34,586; mean: 629 reads; SD: 1289) in the negative controls (NC; *n* = 58). In these NC, 90% of the reads represented 11 variants, and 38% belonging to a human haplotype that was detected with a maximum of 3775 reads in the most contaminated negative control (NC_PCR_). T_CC_ thresholds ranged between 0 and 3775 reads, depending on the variant considered. After filtering the dataset using these T_CC_ thresholds, we kept 5,704,150 reads representing 5697 variants (Table 2).

**Table 2.** Objectives and impacts of the data filtering steps on the number of reads, variants and (putative false and true) positive occurrences of variants in the abundance table.

| Filter step | Objectives | Nb of reads | Nb of variants | Nb of positive occurrences: |  |  |  |
| --- | --- | --- | --- | --- | --- | --- | --- |
|  |  |  |  | for 3 to 3<br>PCR replicates | only for 2 to 3<br>PCR replicates | only for 1 to 3<br>PCR replicates | total |
| Raw abundance table | NA | 5,835,359 | 5751 | 4969 | 4092 | 14,374 | 23,435 |
| Threshold T<sub>CC</sub> | Filter out cross-contaminations generated during the pre-sequencing procedures using the maximal number of reads per variant observed in various negative controls | 5,704,150 | 5697 | 4689 | 3496 | 11,618 | 19,803 |
| Threshold T<sub>FA</sub> | Filter out misindexing generated during the sequencing using the rate of read false assignment calculated thanks to a DNA internal control (here a beluga whale DNA) | 5,684,166 | 5697 | 4466 | 3243 | 9527 | 17,236 |
| PCR replicates | Eliminate inconsistent results between the three PCR replicates from the same sample to remove putative false positive occurrences | 5,172,708 | 2636 | 4466 | NA | NA | 4466 |

NA: not applicable

#### Filtering false-assignments of reads using threshold T_FA_

The ‘alien’ positive control (PC_alien_) produced a total of 30,179 reads among which 30,111 were assigned to the nine independent PCRs performed on this DNA. The other 68 reads, i.e. 0.23%, were mis-assigned to 52 other samples, with a maximum of 13 reads observed for a given bat faecal sample. The maximum false assignment rate R_FA_ was therefore equal to 0.043%, and T_FA_ varied between 0 and 244 reads depending on the variant considered. After filtering, the result table included 5697 variants and 5,684,166 reads. Note that T_FA_ excluded reads but not taxa (Table 2).

#### Filtering inconsistent results using the three PCR replicates

74.1% of occurrence (i.e. cells showing at least one read in the abundance table) were not replicated and were removed. Among these inconsistent results, 74.6% were positive for only one of the three PCR replicates. The remaining reads represented 2636 variants and 5,172,708 reads (Table 2).

Finally, for each sample and variant, the reads of the replicated PCRs were summed in the abundance table. The 21 bat samples that did not include any read after data filtering were discarded from the dataset.

### Comparison of DNA extraction kits

We removed the NucleoSpin Soil kit of this comparison regarding the low number of samples (*n*=9). The five other DNA extraction kits differed in their performance levels (Table 3, Table S1). Statistical results are detailed in Appendix S2.

**Table 3.** Comparison of six DNA extraction kits used for the metabarcoding of bat faecal pellets.

| DNA extraction kit | Sample size | Mean number of reads per PCR | PCR failure (<500 reads) | Success of sequencing |  |
| --- | --- | --- | --- | --- | --- |
|  |  |  |  | Host (bat) | Prey (arthropod)<sup>§</sup> |
| Dneasy mericon Food Kit (Qiagen) | 47 | 3489 | 7% | 68% | 83% |
| EZ-10 96 DNA Kit, Animal Samples (BioBasic)<sup>†</sup> | 113 | 4822 | 14% | 70% | 81% |
| NucleoSpin 8 Plant II (Macherey-Nagel)<sup>†</sup> | 95 | 6712 | 1% | 84% | 88% |
| NucleoSpin Soil (Macherey-Nagel)<sup>‡</sup> | 9 | 7300 | 0% | 67% | 89% |
| QIAamp Fast DNA Stool Mini Kit (Qiagen) | 47 | 2901 | 14% | 70% | 79% |
| ZR Fecal DNA MiniPrep (Zymo) | 46 | 6209 | 1% | 72% | 80% |

<sup>†</sup>96-well format for high throughput DNA isolation
<sup>‡</sup>removed of the statistical analysis
<sup>§</sup>including unclassified arthropod that were not recorded in the sequence databases

The total number of reads obtained per sample and the sequencing success of bats were significantly influenced by the extraction kits, but not by storage duration. The NucleoSpin 8 Plant II DNA extraction kit produced the highest number of reads. This variation resulted partly from PCR failures whose rate could reach 14% (QIAamp Fast DNA Stool and EZ-10 96 DNA). The NucleoSpin 8 Plant II DNA extraction kit also lead to the highest success of bat sequencing. 84% of the faecal samples analysed with this kit produced replicable bat identifications (only 68% to 72% of the samples with the four other kits).

The sequencing success of preys did not depend of the extraction kit, although the NucleoSpin 8 Plant II provided the best result. Prey sequencing was marginally influenced by storage duration, with a marked decrease being visible after three months storage at room temperature.

### Mock community analyses

The 19 arthropod DNA extracts from the mock communities were first amplified and sequenced independently in the MiSeq run. The MiSeq sequences produced similar identifications than Sanger sequences when considering the 19 most abundant MiSeq variants (Table 1). These 19 variants were 100% identical to the reference Sanger sequences (see the alignment file of the 19 insect species included in the mock communities deposited in Dryad: https://datadryad.org/resource/doi:10.5061/dryad.kv02g/8Dryad). They represented 79% of the reads for these individual PCRs, confirming the high quality of the MiSeq sequencing. BOLD identification tool enabled to identify specifically 14 of the 19 COI minibarcode sequences. *Forficula lesnei* (MC_1_) and *Acorypha* sp. (MC_2_) were not identified using the public databases BOLD and GENBANK because they were not referenced. Concerning *Monochamus sutor*, *Chrysopa perla* (MC_1_) and *Bactrocera dorsalis* (MC_2_), we obtained equivalent multi-affiliations with two to five candidate taxa matching with phylogenetically close species (*Monochamus sutor* or *M. sartor*; *Chrysopa perla* or *C. intima*; *Bactrocera dorsalis*, *B. invadens*, *B. philippinensis*, *B. musae* or *B. cacuminata*).

Other non-expected variants were detected at low frequencies (mean: 0.46%, min.: 0.02%, max.: 17.12%) and corresponded to heteroplasmy, pseudogenes (NUMTs: Nuclear insertions of mitochondrial sequences) or to parasitoid sequences. Indeed, in *Forficula lesnei* sample, the tachinid parasitoid *Triarthria setipennis* was detected (Table 1).

We next analysed results of both mock communities amplified and sequenced in pools. MC_1_ sequencing revealed 11 of the 12 insect expected sequences (Table 1). Frequencies of reads varied from 0.4% to 30.1% (expected frequencies 8.3%) while genomic DNA extracts were mixed in equimolar concentrations, therefore revealing biases in PCR amplification. *Protaetia morio* was not detected after data filtering. Insights into the raw dataset showed that some reads were obtained in all three PCRs but with numbers below T_CC_ threshold (T_CC_ = 5 reads for this variant). Sequences of the parasitoid fly *Triarthria setipennis* were detected at low frequency (0.2%). The seven insect species included in MC_2_ were detected, with frequencies ranging from 1.5% to 27.3% (expected frequencies 14.3%). For both mock communities, chimeric reads were detected visually despite the filtering processes using UCHIME program, with frequency levels reaching 3.4% and 5.1%, for MC_1_ and MC_2_ respectively (see the data for the mock communities in the abundance table after filtering deposited in Dryad: https://datadryad.org/resource/doi:10.5061/dryad.kv02g/8).

### Taxonomic identification of bats and their preys

Up to now, less than half of COI sequences deposited in BOLD Systems are made public (Species Level Barcode Records in January 2017: 2,697,359 Sequences/175,125 Species; Public Record Barcode Database: 1,018,338 Sequences/85,514 Species). Because we could not identify all variants using the web application of BOLD Systems, we decided to analyse the 1318 most abundant variants of the whole dataset (including bat samples and mock communities). They were represented by more than 100 reads what was equivalent to 99.1% of all remaining reads. Further analyses revealed that a majority of the remaining 1318 rare sequences (less than 100 reads) could not be assigned clearly to any taxa. They are mainly chimeric sequences or pseudogenes that had not been removed during the filtering process (see the data for the mock communities in the abundance table after filtering deposited in Dryad: https://datadryad.org/resource/doi:10.5061/dryad.kv02g/8).

The analysis of the 336 bat faecal samples in three PCR replicates led to the detection of 1080 abundant variants and 1232 rare ones. The abundant variants included 47 variants assigned to bat species (1,974,394 reads) and 925 variants belonging to the phylum Arthropoda (1,619,773 reads). Among these latter, 654 variants were assigned with similarity level higher than 97% in BOLD (1,305,633 reads). Finally, 35 variants could not be assigned to any taxa either in BOLD or GENBANK (80,977 reads) (Fig. 2 and Fig. S2). Within samples, the proportion of reads between bats and arthropods was quite balanced, except for *Myotis nattereri*, *Myotis mystacinus* and *Myotis alcathoe* for which lower frequencies of reads from bats were observed (Fig. S2).

**Figure 2.**
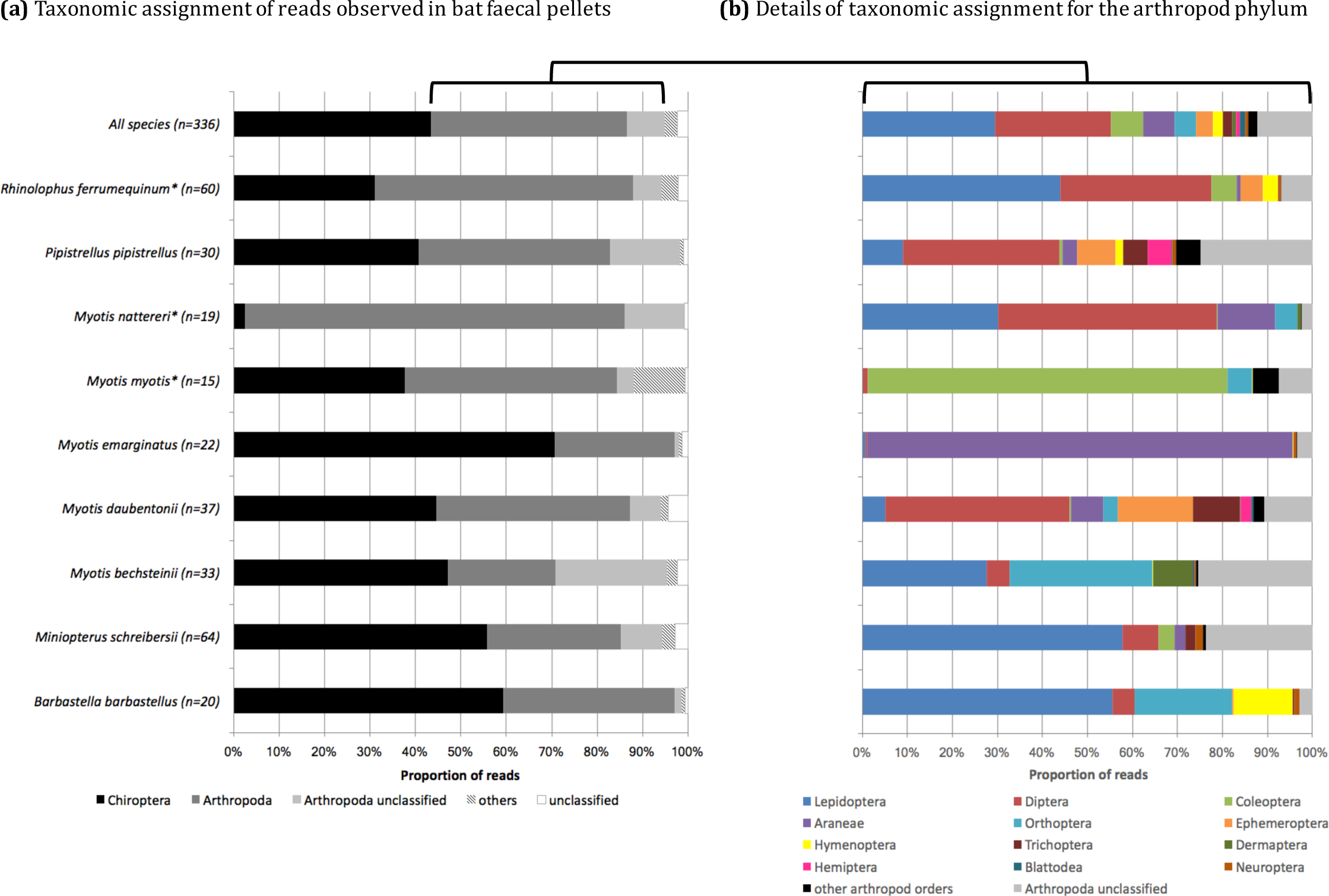
Proportion and taxonomic assignment of reads obtained from high-throughput sequencing of bat faecal pellets. The “All species” bars describe results obtained for all 336 samples from the 16 bat species. Specific results are only detailed for bat species whose sample sizes were over 19. (a) Histogram showing the proportion and assignment of all reads. “Others” (striped bars) includes reads that corresponds to sequences that likely belong to organisms from blood meal/coprophagia/necrophagia or secondary preys, parasites, diet of insect and putative contaminants. (b) Histogram detailing the proportion and assignment at the order level of reads corresponding to arthropod phylum only. * indicates bats species showing a substitution T⤍C that creates a mismatch between the COI target sequence and the position 3’-end of the reverse PCR primer.

Surprisingly, other non-expected taxa were also detected, including 73 variants (133,423 reads) attributed to nematodes (39,186 reads), plants (28,444 reads), gastropods (19,915 reads), algae or fungi (18,972 reads), rotifers (11,925 reads), tardigrades (462 reads), birds (285 reads) as well as mammals (14,234 reads). The 1232 remaining rare variants (32,868 reads) corresponded to 0.7% of the 4,932,226 reads and were considered as unclassified (Fig. 2, Fig. S2).

### Comparison of field and molecular identification of bat species

As expected, the COI minibarcode was resolutive to the species level, except for two species sampled, *Myotis myotis* and *Eptesicus serotinus*. Their assignments were equivalent between two species, *M. myotis* and *M. blythii* in one hand, and *E. serotinus* and *E. nilssonii* in the other hand. We found a congruent taxonomic identification of bat species between molecular and morphological analyses for 238 out of the 336 faecal samples analysed (70.8%). The 98 remaining samples that did not provide a reliable taxonomic identification mainly resulted from amplification failures for at least one PCR replicate (72 samples, i.e. 21.5%: for one (5.4%), two (5.4%) or the three (10.7%) PCR triplicates performed for each sample). Details are provided in Fig. S3. They mostly concerned *Myotis nattereri* (12 failures over 19 samples tested) and *Rhinolophus ferrumequinum* (24 failures over 60 samples tested). A mismatch (T/C) at the 3’-end of the reverse primer could be at the origin of these higher rates of amplification failure for these bat species (Table S4). Finally, 17 samples provided ambiguous molecular results with two bat species detected for one pellet and ten samples lead to incongruent taxonomic identification with regard to bat morphology (Fig. S3).

### Diet composition

In further diet analyses, we considered the 268 bat faecal samples for which we had a congruent taxonomic identification between morphological and molecular analyses based on the results of one (*n* = 15 samples), two (*n* = 15) or three (*n* = 238) PCR replicates (Fig. S3). We removed samples for which no prey data could be analysed, including 20 samples for which only bat sequences were recovered, six samples with only unclassified sequences, eight samples with only sequences of nematodes, plants, fungi and/or algae, and 18 samples for which arthropod sequences were recovered, but with levels of similarity that were too low (<97%) to provide a reliable assignation to a precise taxon. Altogether, diet compositions were described on 216 bat faecal samples, corresponding to 16 bat species. Among the 551 validated arthropod variants of these samples, we identified 18 arthropod orders, 117 family, 282 genus and 290 species (Table S5). We observed a wide heterogeneity in the taxonomic diversity and composition of diets between bat species (Fig. 3).

**Figure 3.**
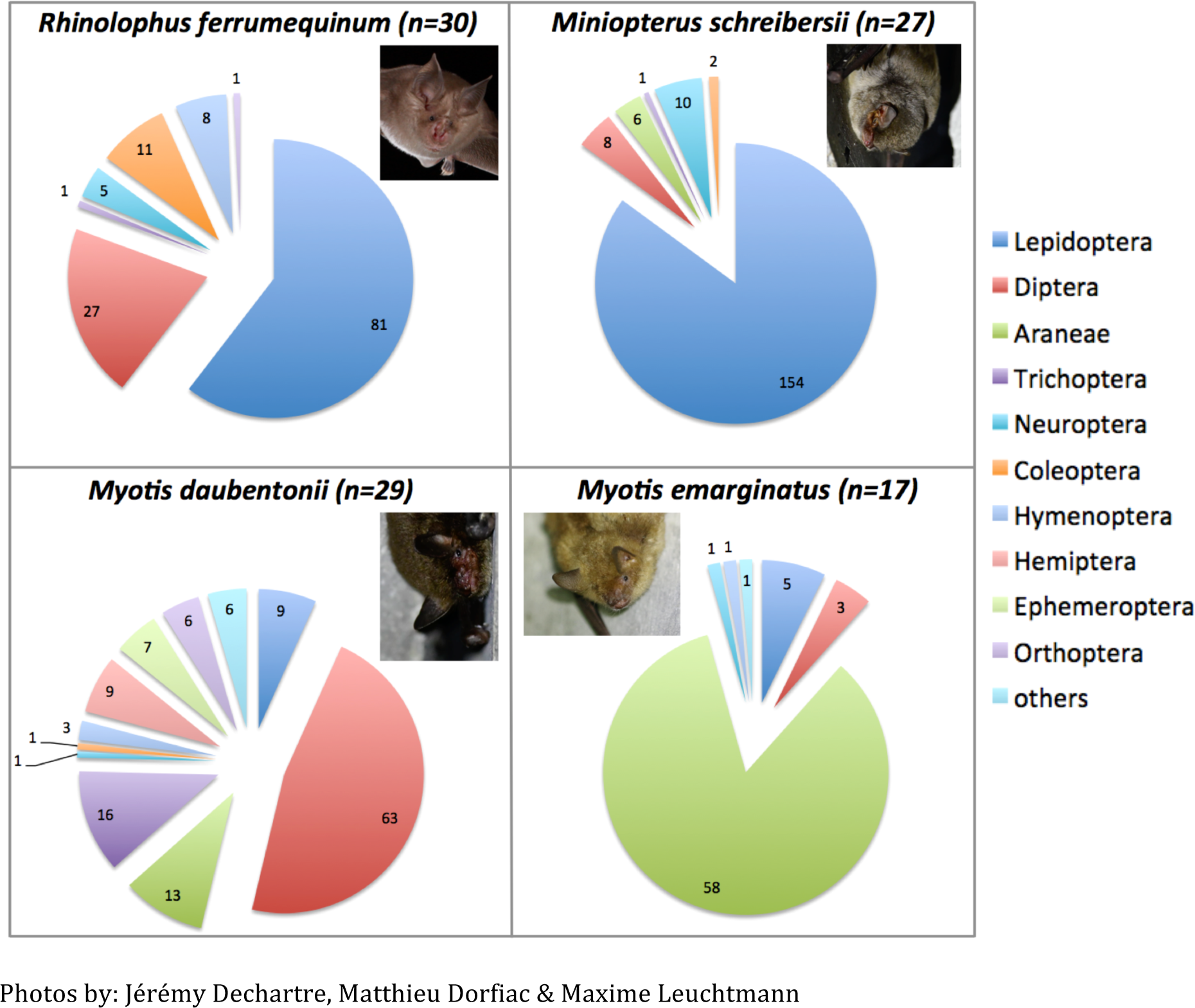
Number of occurrence of prey taxa within faecal pellets of four bat species with contrasted diets. The details of taxa names and occurrences for the diet of the 16 bat species are reported in Table S5 (Supplemental Information).

## Discussion

### Importance of data filtering and controls

Assessing diet variability between individuals and populations is of main importance to better understand the biology of species, here bats. Although the procedures limiting false positive results have been well described (Ficetola et al., 2016), they still remain scarcely included in methodological procedures (but see in an epidemiological context, Galan et al., 2016). Negative controls are often included during DNA extraction and PCRs, they are not always sequenced and only checked using gel electrophoresis. This procedure is not satisfactory as most contaminating sequences cannot be visually detected. In particular, cross-contaminations of index/tags, tags jumps (Schnell et al., 2015) or polyclonal / mixed clusters during Illumina HTS (Kircher et al., 2012) that may lead to the mis-assignments of reads. The positive control PC_alien_ proposed here enabled to estimate these read mis-assignments for the whole run (0.23%) or for a given sample (up to 0.043%).

Here, we proposed a set of filtering and validation procedures based on negative, positive controls and three independent PCR replicates. This strategy based on non-arbitrary filtering thresholds has recently been applied in Corse et al. (2017) to study the diet of a critically endangered fish species. Our results showed that applying T_CC_ and T_FA_ thresholds removed relatively few reads and variants compared to the replicate validation procedure. Hence the rates of laboratory contaminations or mis-assignments during HTS seemed to be low while the proportion of non-repeatable variants seemed to be high. The reasons might be methodological (e.g. PCR chimera, sequencing errors or PCR drop-out) or biological (presence of NUMTs at low frequencies, low biomass preys, traces of ancient meals). Despite our filtering pipeline, an important proportion of variants that were validated by the three replicates remained at low frequencies. They can be attributed to chimera, pseudogenes or PCR and non-random sequencing errors (see the data for the mock communities in the abundance table after filtering deposited in Dryad: https://datadryad.org/resource/doi:10.5061/dryad.kv02g/8). We therefore had to eliminate 50% of the less frequent variants (1318 variants over 2636) in order to make taxonomic assignment in BOLD Systems easier. This important number of potentially artefactual variants could be decreased by applying the filtering procedure recently proposed by Corse et al. (2017). In addition to the non-arbitrary filters described in our method, they applied algorithms that removed errors (Obiclean, see Boyer et al., 2016), filtered chimera more efficiently (UCHIME 2, Edgar, 2016) as well as pseudogenes. These procedures enabled to keep a high proportion of reads (70%) and a low proportion of variants (0.3% corresponding to 61 to 81 variants depending on the minibarcodes considered). The cross-comparison of taxonomic identification results obtained with different assignment methods is therefore feasible for the variants validated (Corse et al., 2017). Finally, the sensibility of prey detection could be improved by applying the relaxed restrictive approach of between-PCR replicate validation proposed by Alberdi et al. (2017). As such, variants found in at least two of the three replicates could be considered positive, but this strategy may also increase false positive results.

Finally, mock communities are not systematically tested in animal metabarcoding studies (but see Pinol et al., 2015) although it enables to empirically assess the efficiency and biases of both molecular designs and bioinformatic pipelines. We recommend to include diversified artificial communities in metabarcoding studies, ideally encompassing the whole potential phylogenetic diversity of the samples to be studied.

### Methodological framework to avoid biases

Fieldwork remains a crucial step for diet analyses, even with such molecular approaches. The way faecal pellets are collected may lead to cross-individual contaminations, as revealed in our study by the detection of bat sequences corresponding to the wrong species. Considering invasive sampling, cotton holding bags in which individuals are kept before morphological analyses must be carefully checked to avoid the collection of faecal samples belonging to successively captured bats. When possible, a disposable collection system or a UV decontamination procedure between captures could be performed. Faecal pellets should also be carefully handled and stored to avoid contaminations. These points reinforce the importance of performing species bat identification for each pellet to prevent mis-assignment of a diet and to an individual of the wrong species. In the case of non-invasive sampling, it is highly recommended to use clean supports and single-use instruments to collect the pellets. Reducing time-delay between bat faeces release and collection might also be important to avoid DNA degradation (Oehm, Juen, Nagiller, Neuhauser, & Traugott, 2011) and cross-contaminations due to urine from different species of bats or coprophagous arthropods for example.

Further storage conditions of faecal samples are of main importance to guarantee DNA integrity and limit the proliferation of micro-organisms. The rate of amplification failures observed in our study were hence likely to result from the storage of all the bat faecal samples at room temperature in empty tubes during several months. Samples should be frozen at −20°C or in liquid nitrogen, or stored in an appropriate storage buffer (e.g. ethanol) to prevent DNA degradation (Renshaw, Olds, Jerde, McVeigh, & Lodge, 2015). The method proposed here enabled to obtain satisfying results even if samples had not been stored in the optimal conditions required for metabarcoding analyses. It is therefore likely that samples that were not initially collected for genetic/metabarcoding purposes, potentially including ancient samples (from guano collected several weeks after dropping to guano accumulated during decades), could also be successfully assessed for diet analyses using our sequencing protocol.

We also showed that extraction methods may influence the success of metabarcoding studies. Our results evidenced that NucleoSpin 8 Plant II kit provided the best results in terms of host sequencing success, number of reads produced per PCR and prey sequencing success. It therefore seems to be the best compromise between cost per extraction, throughput (96-well format) and quality of the DNA purification for metabarcoding applications.

In addition, we have evidenced PCR amplification biases for particular prey and bat taxa that lead to a high variability in sequencing depth and even to the amplification failure for particular species. Sources of PCR biases can be multiple (too stringent PCR conditions (e.g. high hybridization temperature), differential DNA degradation, interspecific mitochondrial copy number variation…). Nevertheless primer mismatches are one of the most important source of PCR biases (Pinol et al., 2015). A *post-hoc* analysis performed on 693 COI sequences available for the 33 bat species found in France revealed that 17 bat species had a frequent mismatch at the 3’-end of the reverse primer (see Table S4). The analyses of mock communities (MC) also revealed amplification biases with regard to arthropod species, as previously described in other empirical studies (Pinol et al., 2015). It was probably due to the same 3’-end mismatch described above between PCR reverse primer and arthropod species DNA, and to further primer annealing competition during DNA amplification of the community. Indeed, the four species that showed the lowest proportion of reads (*Calliptamus barbarus*; *Phymata crassipes*; *Protaetia morio*; MC_2_: *Bactrocera dorsalis*, see Table 1) have this mismatch in their sequences. For future studies, we recommend a new version of the target-specific reverse primer (modifications in bold: MG-univR-MiSeq 5’-ACTATAAA**RA**A**R**A**TY**ATDAY**R**AADGCRTG-3’) to limit the observed biases for most bat and arthropod species.

Recent studies have also proposed the use of several minibarcoding primer sets (Alberdi et al., 2017; Corse et al., 2017) to maximize the taxonomic coverage of metabarcoding approaches and minimize false negative results. Indeed, it is noteworthy that designing COI universal primers generating no PCR amplification biases might not be achievable (Deagle, Jarman, Coissac, Pompanon, & Taberlet, 2014). Therefore the use of ribosomal rRNA, either mitochondrial 16S or 12S, with conserved flanking regions among taxonomical distant species, might be encouraged. Although this proposal should reduce amplification biases among taxa, it is yet not suitable for arthropod metabarcoding approaches due to the absence of public databases similar to BOLD (i.e. curated database with reference sequences linked to taxonomically verified voucher specimens). Moreover, the taxonomic resolution of these barcodes at the species level remaining largely unknown (Elbrecht et al., 2016). In conclusion, COI minibarcodes still remain an imperfect, but ‘not so bad’ solution for taxonomic identification when this latter requires to reach the species level.

### Prey-bat simultaneous identification and taxonomic resolution

In addition to the methodological framework provided, the originality of our approach also resided in the simultaneous identification of preys and of a wide diversity of bat species. Despite the low quality of our faecal samples due to inappropriate storage conditions, we obtained a specific identification for 74% of the bat samples studied, including only 3% of incongruent results between the molecular and field identifications due to remaining faecal pellets in the cotton holding bags (see above). Among the 26% unidentified, 21% were due to bat sequencing failure of at least one PCR replicate, 5% corresponded to the detection of several bat species. The failure rate associated with our molecular approach was therefore equivalent with what is observed in studies where species identification from bat faecal pellets was performed using traditional Sanger sequencing (e.g. 19% reported in Hope et al., 2014). It could be easily improved with the use of appropriate storage conditions and DNA extraction method. Our approach is thus relevant for bat species identification too.

The simultaneous identification of predators and preys is generally avoided in diet analyses as it may induce biases in the pool of sequences produced (Pompanon et al., 2012). In particular, high success of predator amplification will reduce prey amplification, what will in turn affect the sensibility of diet analyses. Here, we reported well-balanced proportions of reads for bats and their preys, with a mean of 43.3% of bat reads whereas other studies reported higher proportion of predator sequences (91.6% for the leopard cat in Shehzad et al., 2012). The lower proportion of bat reads observed in our study may result from several phenomena: a lower proportion of predator DNA in the faecal pellets analysed, a lower DNA degradation of prey due to the very quick bat digestion in bats, or a lower primer specificity to amplify target DNA from bats. It enabled an important sequencing depth for arthropods that guarantees the high sensitivity for prey detection. However, few samples produced low numbers of arthropod sequences. Results between PCR replicates were therefore not repeatable, what led to potential false negatives. The increase of sequencing depth, for example using an Illumina HiSeq platform, associated with the relaxed restrictive approach of between-PCR replicate validation (see above) could improve our ability to detect preys.

Our approach provided a high level of taxonomic resolution for bats (Fig. S1) and their preys as evidenced by the congruent identifications obtained using the 658bp Sanger sequences and the 133bp minibarcode for the arthropod mock community. Particularly, we confirmed the possibility to discriminate morphologically close insect species including the pine processionary moth complex *Thaumetopoea pityocampa*/*T. wilkinsoni* (Kerdelhue et al., 2009), the longhorn beetle *Monochamus galloprovincialis* and its sister species *M. sutor* (Haran, Koutroumpa, Magnoux, Roques, & Roux, 2015) or the green lacewing *Chrysopa perla* / *C. formosa* (Bozsik, 1992). Such taxonomic resolution is highly important when dealing with arthropod pests or arthropod species involved in biological pest control. In particular cases (Mayer & von Helversen, 2001), knowledge on geographic distribution may help deciphering the most likely taxa in presence (e.g. *Eptesicus serotinus* and *Eptesicus nilssonii*, or *Myotis myotis* and *Myotis blythii* that cannot be distinguished whatever the mitochondrial marker used).

We have also evidenced some limitations with regard to public sequence databases. Two arthropod species included in the mock communities could not be identified as they were not included in BOLD and G_EN_B_ANK_ databases. We have also emphasized some errors that may have consequences for further identification (Fig. S1). For example, one sequence of *Pipistrellus pipistrellus* included in BOLD is mis-assigned to *Pipistrellus kuhlii* (G_EN_B_ANK_ Accession JX008080 / BOLD Sequence ID: SKBPA621-11.COI-5P). This mistake made it impossible to distinguish both species using BOLD Systems tool. Although recently reported (Shen, Chen, & Murphy, 2013), it has not yet been cleaned. It seems that such errors are quite frequent in G_EN_B_ANK_ (Shen et al., 2013), and unfortunately BOLD database does not appear to be spared. Completeness and reliability of public databases are therefore still a main pitfall in metabarcoding studies.

### Dietary composition for 16 bat species in Western France

The combination of HTS and filtering procedures described here provided a detailed diet characterization of the 16 bat species sampled in Western France. At the order level, our results were congruent with previous knowledge of preys consumed detected using morphological analyses. For example, the diet of *Myotis daubentonii* and *Pipistrellus pipistrellus* is known to be dominated by Diptera (e.g. Arlettaz, Godat, & Meyer, 2000; Vesterinen, Lilley, Laine, & Wahlberg, 2013), what was confirmed by our results showing respectively 79% and 74% of faeces samples positive for Diptera. Lepidoptera were also highly frequent in *Pipistrellus pipistrellus* samples, being the second order detected in faecal samples (48% of positive faecal pellets), as described in Arlettaz et al. (2000). The diet of *Barabastella barbastellus* was dominated by Lepidoptera with 100% positive samples, as previously described in Andreas et al. (2012). Finally, the lowest dietary richness was found for *Myotis emarginatus*, with sequences of Arenae detected in 100% of faecal samples, what was previously described in Goiti et al. (2011).

Compared to morphological studies that enabled prey taxonomic identification at the order (sometimes family) level (Arlettaz et al., 2000), our approach provided greater details on dietary composition by increasing prey taxonomic resolution. It may even allow distinguishing species that could not be recognized based on morphological criteria of mixed insect hard parts. In addition, our approach enabled to detect unexpected interactions including secondary predation events and gastrointestinal infestations. We reported the presence of snails (*Cepaea hortensis* and *Cepaea nemoralis*) and slugs (*Arion intermedius*) in diets that included Carabidae preys (*Abax parallelepipedus* and *Carabus* sp.) in three *Myotis myotis* samples. Also surprisingly, three *Myotis nattereri* and one *Myotis daubentonii* bat faecal samples contained *Bos taurus* sequences. It is likely that these findings result from traces of bovine animal blood in the gut of the biting house fly *Stomoxys calcitrans* that were also detected as consumed preys in the *M. daubentonii* sample, or traces of bovine animal excrements coming from green bottle fly (*Neomyia cornicina*), crane flies (*Tipula* sp.) or scavenger cokroach (*Ectobius* sp.) for the *M. nattereri* samples. Similarly, grey heron (*Ardea cinerea*) and dormouse (*Glis glis*) sequences found in two *Myotis daubentonii* bat faecal samples may be due to blood traces in biting insects, the common house mosquito *Culex pipiens* and the autumn house-fly *Musca autumnalis* respectively. These results must be considered with caution because field contamination cannot be fully discarded. In addition, secondary predation can sometimes be difficult to distinguish from primary predation. Nevertheless the detection of prey diet traces in bat faeces indicates the possibility to use our results to improve our knowledge of trophic relationships.

Altogether, our dataset enabled to reveal the presence in bat diets of 61 pest species (Table S5) among which some are important for agricultural management (e.g. the cotton bollworm *Helicoverpa armigera*, the spotted-wing drosophila *Drosophila suzukii* and the pine processionary *Thaumetopoea pityocampa*) or veterinary and Public Health issues (e.g. the malaria vector *Anopheles claviger,* the biting house fly *Stomoxys calcitrans*, the common house mosquito *Culex pipiens*). Our results therefore confirmed the possibility to use our metabarcoding approach as an indirect tool for “chirosurveillance” without any *a priori* with regard to the pests that need to be surveyed (Maslo et al., 2017). This study also illustrated the capacity of our approach to reveal variation in diet richness and composition. Combining bat-species molecular identification with diet analyses will provide a more complete understanding of how bat diet varies along season, life history stage, gender and age. It will be of main importance to understand the influence of diet on bat fitness and colony viability and to answer questions about niche size and niche overlap for co-existing species.

## Conclusion

The DNA metabarcoding approach described here enables the simultaneous identification of bat species and their arthropod diets from faeces, for several hundreds of faecal samples analysed at once. This strategy reduces the number of molecular steps than usually required in other metabarcoding studies and minimizes the probability to mis-assigned preys to the wrong bat species. The two-step PCR protocol proposed here makes easier the construction of libraries, multiplexing and HTS, at a reduced cost (about 8€ per faecal pellet for the entire wet lab workflow). Our study also includes several controls during the lab procedures associated to a bioinformatic strategy that enables to filter data in a way that limits the risk of false positive and that guarantees high confidence results for both prey occurrence and bat species assignment. This study therefore provides a rapid, resolutive and cost-effective screening tool for addressing ‘chirosurveillance’ application or evolutionary ecological issues in particular in the context of bat conservation biology. It may be easily adapted for use in other vertebrate insectivores, and more widely for other amplicon sequencing applications.

## Acknowledgements

First of all, we are indebted to all of our collaborators in the naturalist associations who helped us to gather and generate this comprehensive bat sampling: Matthieu Dorfiac (Charente Nature), Anthony Le Guen (Deux-Sèvres Nature Environnement), Virginie Barret and Philippe Jourde (LPO France), Sébastien Roué (Groupe Chiroptères Aquitaine) and all the numerous volunteers that contributed to field work.

We thank Jean-Claude Streito, Guenaelle Genson, Antoine Foucard, Laure Benoit, Carole Kerdelhué and Laure Sauné for the insect DNA provided to constitute the mock communities. Many thanks to Emese Meglecz, Vincent Dubut and Emmanuel Corse for the constructive discussions and sharing about the metabarcoding analyses. We thank Eric Petit and two anonymous reviewers for their constructive comments and suggestions on a previous version of the manuscript. Data used in this work were partly produced through the genotyping and sequencing facilities of ISEM (Institut des Sciences de l’Evolution-Montpellier) and LabEx Centre Méditerranéen Environnement Biodiversité. Analyses were performed on the CBGP HPC computational platform thanks to Alexandre Dehne-Garcia. This work was funded by the LabEx ECOFECT and the internal funding from the CBGP laboratory. Orianne Tournayre PhD is funded by the LabEx CeMEB. Part of this work (JBP, DP) was performed within the framework of the LabEx ECOFECT (ANR-11-LABX-0048) of Université de Lyon and the Poitou-Charentes Nature Feder project « *Grand Rhinolophe et trame verte bocagère: étude des facteurs environnementaux influant sur la dynamique des populations* » (ML, JBP, DP).

## Data accessibility

Supplementary data deposited in Dryad (under embargo during the process of peer review) include: i) raw sequence reads (fastq format), ii) raw output files generated by the M_OTHUR_ program, iii) raw abundance table, iv) filtered abundance table including taxonomic affiliations, v) alignment of 444 COI minibarcode sequences from BOLD corresponding to 33 bat species found in France used to construct the phylogenetic tree, vi) alignment of 693 COI reverse primer target sequences from BOLD corresponding to 33 bat species found in France and vii) alignment of the COI haplotypes obtained by Sanger and MiSeq sequencing for the 19 insect species included in the mock communities. Supplementary data and information are available on request to the corresponding author.

## Author contributions

The study was conceived and designed by M.G., D.P. and N.C. J.-B.P and M.L. supervised the field work. M.G. and J.-B.P carried out the molecular biology procedures and validated the MiSeq data. M.G. contributed to the development of bioinformatics methods and M.G., J.-B.P and E.P. validated taxonomic assignments. D.P. and N.C. coordinated the funding projects (resp. Ecofect and AAP CBGP). M.G., J.-B.P, O.T. and N.C. analysed the data. M.G. and N.C. wrote the manuscript. J.-B.P, O.T., E.P., M.L and D.P. helped to draft and to improve the manuscript. All coauthors read and approved the final manuscript.

## Supporting Information

**Table S1:** Information about the samples, the laboratory controls and the technical replicates

**Table S2:** Taxonomic resolution of the 133bp COI minibarcode. A phylogenetic tree is built on the basis of 444 COI reference sequences corresponding to 33 bat species found in France. Bold sequences ≥ 132bp for the minibarcode and without N were selected.

**Table S3:** Objectives and impacts of mothur program steps on the number of reads and variants

**Table S4:** Mismatches between the COI reverse primer used in this study and 693 reference sequences corresponding to 33 bat species found in France

**Table S5:** Table S5 List of preys identified within faecal pellets for 16 bat species using high-throughput sequencing. Only results with sequence similarity >97% are kept. We applied a modified version of the criteria described in Razgour et al. (2011) for the confidence level of the sequence assignments: 1a: match to only one species in BOLD System and >99% sequence similarity; 1b: match to only one species in BOLD System and >98% sequence similarity; 2: several species of the same genus and >98% sequence similarity; 3: several species of different genus of the same family and >98% sequence similarity; 4: several species of different families and/or >97% to <98% sequence similarity. See the different sheets of this file for the details per bat species

**Figure S1:** Neighbor-Joining tree obtained from the analysis of the 133bp minibarcode (COI) of bat species present in France (444 sequences, 33 species). The evolutionary distances were computed using Kimura 2-parameter method. Tree robustness was assessed using a bootstrap with 500 replications. Species names in red indicate the pairs of bat species that could not be discriminated because they were assigned to the same genetic cluster. The specific identifications of samples in red squares may not be reliable due to taxonomic mis-identification or weak quality of sequencing.

**Figure S2:** Proportion and taxonomic assignment of reads obtained from high-throughput sequencing of faecal pellets from 16 bat species. Chiroptera reads (black bars) and Arthropoda reads (dark grey bars) correspond to sequences assigned to one or several species using BOLD or G_EN_B_ANK_ and identity scores higher than 97%. “Arthropoda unclassified” reads (light grey bars) correspond to sequences assigned to Arthropoda phylum with identity scores lower than 97%. “Others” (striped bars) includes sequences from putative contaminants (i.e. human, cat or fungi), blood meal/coprophagia/necrophagia (others mammalia or birds), secondary preys (i.e. Mollusca), parasites (Nematoda) and insect diets (i.e. plants). “Unclassified” (white bars) corresponds to unique sequences with no match in BOLD and G_EN_B_ANK_. The “All species” bars describe results obtained for all 336 samples from the 16 bat species.

*indicates bats species showing a substitution T⤍C that creates a mismatch between the COI target sequence and the position 3’-end of the reverse PCR primer.

**Figure S3:** Proportion of faecal pellets with true or false bat species molecular identification using high-throughput sequencing. The “All species” bar describes results obtained for all 336 samples from the 16 bat species. Black, dark grey and light grey bars show the proportion of samples for which morphological and molecular identifications were identical for respectively three, two or one PCR replicate. White bars indicate the proportion of samples for which molecular identification failed for all three replicates. Striped bars show the proportion of samples which morphological differed from molecular identifications, whatever the number of PCR replicates concerned. Dotted bars indicate the proportion of samples for which molecular identification indicated two different bat species names. These latter are likely to be due to field contaminations.

*indicates bats species showing a substitution T⤍C that creates a mismatch between the COI target sequence and the position 3’-end of the reverse PCR primer.

**Appendix S1** Definition and use of negative and positive controls included in the metabarcoding experiment.

**Appendix S2** Factors shaping DNA extraction kit efficiency (number of reads, sequencing success with regard to bats and preys) evidenced by generalized linear models. *P-value* was considered significant when < 0.05.

**Appendix S3** Taxonomic assignment strategy

